# Host dietary specialization and neutral assembly shape gut bacterial communities of wild dragonflies

**DOI:** 10.1101/357996

**Authors:** Rittik Deb, Ashwin Nair, Deepa Agashe

## Abstract

Host-associated gut microbial communities can have large impacts on host ecology and evolution, and are typically shaped by host taxonomy and diet. Different host species often harbor distinct microbial communities, potentially because (1) host dietary specialization determines microbial colonization, (2) host-specific selection acts on diet-acquired microbiota, and (3) a combination of both processes. While the first possibility involves passive community structuring, the other two may arise from a functional association and should produce stable microbial communities. However, these alternatives have rarely been tested in wild host populations. We used 16S rRNA amplicon sequencing to characterize the gut bacterial communities of six dragonfly species collected across multiple seasons and locations. We found that variation in bacterial community composition was predominantly explained by sampling season and location, and secondarily by host species. To distinguish the role of host dietary specialization and host-imposed selection, we used insect-specific primers to identify prey in the gut contents of three focal dragonfly species. We found that these dragonflies – considered to be generalist predators – consumed distinct prey, with seasonal diet variation. Together, the patterns of host dietary specialization and spatial and temporal variation suggest a strong role of passive processes in shaping the gut bacterial community. Indeed, the abundance and distribution of ~76% of the bacterial community members were consistent with neutral community assembly. Our results contradict the pervasive expectation that host-imposed selection shapes gut microbial communities, and highlight the importance of joint analyses of variation in host diet and gut microbial communities of natural host populations.

## 1. INTRODUCTION

Extensive research in the past decade suggests that host-associated gut microbial communities can have large impacts on host evolution (Dillon & Dillon 2004; McFall-Ngai *et al.* 2013; Engel & Moran 2013). Hence, many studies have tried to understand the processes that determine the composition of gut microbiota (Dillon & Dillon 2004; Engel & Moran 2013). It is clear that the gut microbiome is affected by multiple factors including host genotype, environmental variation, and host diet. For example, in mice, knocking out single host genes had a remarkable effect on the gut microbial composition (as reviewed in Spor *et al.* 2011). Even between genetically closely related hosts, environmental variation can create a significant deviation in gut microbiota. For instance, when reared in distinct environments, pairs of human twins (Zoetendal & Akkermans 2009; Nelson 2011; Kostic *et al.* 2013), mice siblings (Gootenberg & Turnbaugh 2011), and *Drosophila* populations reared on the same diet – all harbored distinct gut microbiota (Dillon & Dillon 2004; Charroux & Royet 2012; Broderick & Lemaitre 2012; Engel & Moran 2013). Similarly, variation in host diet may also have a large impact on gut microbial composition (Engel & Moran 2013), as observed in laboratory populations of mice, bees and flies maintained in an otherwise constant environment (Ley *et al.* 2008; Turnbaugh *et al.* 2009; Sharon *et al.* 2011; Sullam *et al.* 2012; Moreira *et al.* 2012; Colman *et al.* 2012; Scott *et al.* 2013). However, it is not always clear whether these effects of host genotype, diet and environment reflect variation in the acquisition or the establishment step of microbial community assembly.

In general, gut microbes are acquired from the mother or through the diet; and they may either colonize and proliferate in the gut, or fail to establish. At each step, various stochastic vs. deterministic, and neutral vs. selective processes determine community composition. For instance, a host may consistently acquire a specific set of microbes when they are maternally transmitted, or if the host is a dietary specialist. Within the host gut, microbial survival and growth dynamics may then be determined largely by stochastic neutral processes (e.g. based on initial abundance); or by deterministic and selective processes such as interactions with the host or with other microbes. Dietary specialists can maintain a specific gut microbial community by constantly reintroducing particular microbes, promoting specific metabolism, and maintaining a consistent gut environment (De Filippo *et al.* 2010; Nicholson *et al.* 2012). In contrast, a generalist host is more likely to stochastically sample a wider range of environmental microbes associated with its variable diet. For example, scavengers and omnivores tend to have richer gut communities (Yun *et al.* 2014; Yadav *et al.* 2015; Shukla *et al.* 2016); and we expect to find a positive correlation between host diet diversity and gut microbial diversity (Engel & Moran 2013; Yun *et al.* 2014). Given this disruptive effect of dietary variation, strong host-imposed selection should stabilize gut bacterial community composition, and minimize the impact of stochastic or deterministic events. Many prior studies have implicated host-imposed selection as a dominant force driving gut bacterial community composition (Spor *et al.* 2011; Engel & Moran 2013; Antwis *et al.* 2017). For instance, host immune responses (Ley *et al.* 2006; Charroux & Royet 2012; Broderick & Lemaitre 2012; Quigley 2013) or a host-derived protected niche inside gut crypts (Dillon & Dillon 2004; Kikuchi *et al.* 2007; Engel & Moran 2013) can selectively allow only specific microbes to colonize the gut. Under weak selection, neutral processes such as ecological drift and microbial dispersal may strongly drive community assembly (Hubbell 2001; Rosindell *et al.* 2011), with each host’s microbiota functioning as a local community interacting with the larger meta-community outside the host body (Costello *et al.* 2009, 2012).

Although the assembly and composition of gut bacterial communities is likely affected by all the processes described above, their relative importance in determining the composition and stability of gut bacterial communities of natural animal populations remains unclear. This gap in our understanding arises partly because most animal studies have focused on genetically homogeneous host populations reared on simple diets in controlled environments, such as laboratories or greenhouses (Engel & Moran 2013) (but see Corby-Harris *et al.* 2007; Osei-Poku *et al.* 2012; Adair *et al.* 2018). In contrast, in nature, most animals occupy diverse spatially and temporally separated niches, with substantial variation in the environment, genetics, behavior, and diet. All these factors can increase stochastic and/or deterministic variation in gut microbiota (Corby-Harris *et al.* 2007; Colman *et al.* 2012; Basset *et al.* 2012; Yun *et al.* 2014), opposing the stability introduced by host-imposed filters. Hence, it is important to ask whether host gut-microbe associations are truly stable in natural populations, and what factors determine the stability and composition of gut microbial communities.

In this context, we analyzed the gut bacterial and dietary community composition in natural populations of six dragonfly species, sampled from six different locations in India, across six months (three seasons) (Fig 1A, and Table S1). Dragonflies are generalist predators of aquatic and associated terrestrial ecosystems (Corbet 2004), and we, therefore, expected that they would consume diverse insect prey across locations, season, and host species. In turn, this dietary diversity should be associated with diverse gut microbial communities. Using models of prokaryotic community assembly, we could specifically test for neutral assembly of communities (Sloan *et al.* 2006) and estimate the proportion of microbes that are assembled neutrally vs. through selection (Chase & Myers 2011). Previously, we found that the culturable fraction of gut bacterial communities of dragonflies varied significantly as a function of host species, location, and sampling time (Nair & Agashe 2016). Here, we built upon this work by sampling more dragonflies, identifying most gut-associated bacteria using 16S amplicon sequencing, and analyzing their diets by amplicon sequencing the cytochrome c oxidase 1 gene (COX1) from gut contents. Using these data, we quantified the spatial and temporal stability of host-associated gut bacteria; tested whether bacterial diversity was correlated with host diet diversity; and quantified the relative importance of neutral processes driving bacterial community assembly.

**Figure 1.**
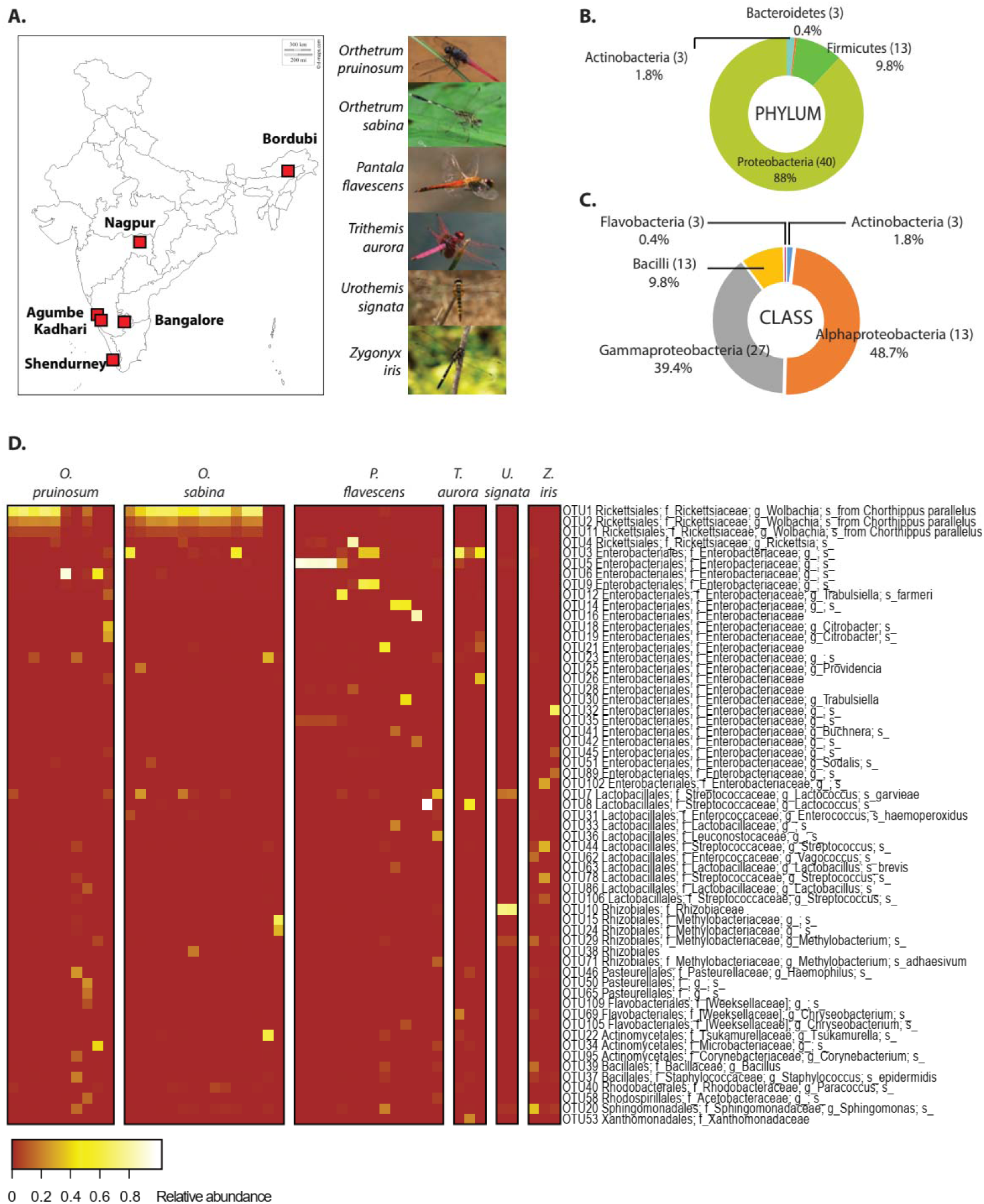
(A) Map of India showing dragonfly sampling locations, and representative images of the sampled dragonfly species. Sampling details are given in Table S1. Major bacterial phyla (B) and classes (C) in the dominant gut bacterial communities of sampled dragonflies. (D) Heat map showing dominant bacterial OTUs across all dragonfly samples. Each column indicates a host individual (sorted by species), and rows indicate dominant bacterial OTUs clustered based on their abundance across hosts.

## 2. MATERIALS AND METHODS

### Sample collection and storage

We collected six dragonfly species from six different sampling sites across India, through three seasons (winter: December – January, summer: March – April, monsoon: October – November) (Fig 1A and Table S1). We caught individuals using butterfly nets in open grounds, near natural water bodies, or waterlogged paddy fields. We conducted three separate rounds of sample collection as follows. (a) To determine the composition of gut bacterial communities, we surface sterilized each dragonfly using 70% alcohol and stored it in a 1×1 ft. mesh cage. Within 4-6 hours of collection, we paralyzed dragonflies using a 4ºC cold shock and dissected them in phosphate-buffered saline (PBS) using sterilized dissection tools. We stored dissected guts in 1.5 ml centrifuge tubes containing 100% molecular grade alcohol. We stored the remaining dragonfly bodies separately in 100% alcohol for subsequent identification using an online resource (http://indiabiodiversity.org/). After bringing samples to the laboratory, we stored them at -20°C until further processing. For collections in Bordubi and Nagpur, we could not dissect dragonflies in the field and so we stored them in 100% alcohol immediately after capture. Note that our sampling was comprehensive across all host species and across locations for 3 host species (*O. pruinosum, O. sabina,* and *P. flavescens*); but we had limited sampling across seasons (Table S1) due to declined population size in dry conditions. (b) To estimate absolute gut bacterial abundance (using qPCR) and to localize bacteria in dragonfly guts (using FISH), we collected and isolated dragonflies in 50 ml Falcon tubes for 4-6 hours so that their guts would be empty. For dragonflies collected for qPCR, we dissected and stored guts in 100% alcohol. For FISH samples, we dissected the gut in PBS, divided each gut into three sections (foregut, midgut, and hindgut), and stored each section separately in 100% alcohol at -20°C. (c) To analyze dragonfly diet, we again collected three of the dragonflies used for gut bacterial community analysis (*Orthetrum pruinosum, Orthetrum sabina* and *Pantala flavescens*) (Fig 1A, Table S1). We isolated individuals in 50 ml Falcon tubes for 4-6 hours to collect fecal matter, and then dissected them to separate gut contents (without host tissue). We stored gut contents and fecal material in separate centrifuge tubes in 100% alcohol at -20°C until further processing.

### Amplicon sequencing to determine gut bacterial and diet composition

We determined the gut bacterial community for a total of 48 dragonflies from different species, geographical locations and sampling seasons (Fig 1A, Table S1). We washed each gut sample thrice in fresh 100% molecular grade alcohol followed by three washes in PBS. We homogenized the tissue in liquid nitrogen using single-use sterile pestles and extracted DNA using the Wizard ^®^ Genomic DNA Purification Kit (Promega Corporations, Wisconsin, Madison, USA). We modified the manufacturer’s protocol as follows: we added 600µl of nuclei lysis solution (10mM EDTA) per 100mg tissue and incubated first at 80°C for 20 min, and then at 65°C for 30 min. We cooled the samples to 55°C, added 20mg/ml proteinase, and again incubated at 55°C for 3 hours. To precipitate degraded protein we added protein precipitation solution and left the sample on ice for 30 min. We centrifuged the lysate at 14000 g for 10 min and precipitated the supernatant with isopropanol. We washed the resulting pellet with 80% alcohol twice, then dried and suspended it in 40µl ultrapure nuclease-free water. We quantified DNA in a Nano-Drop (Nano-drop 2000, Thermo Fisher Scientific Inc., Wilmington, USA) and checked the integrity of the DNA by running 1µg on a 0.8% agarose gel. For each sample, we used 50 ng DNA to PCR-amplify the V3-V4 region of the bacterial 16S rRNA gene, using ExTaq (TaKaRa). The PCR primers contained tag sequences complementary to the Illumina sequencing adapter and index primers from the Nextera XT Index kit V2. We tested amplicons for quality and sequenced them (250 bp paired-end) on the Illumina MiSeq platform (Illumina, San Diego, CA, USA) using standard Illumina forward and reverse primers. Sequencing was performed by Genotypic Technology Pvt. Ltd., Bangalore, India.

For host diet analysis, we implemented a previously described method that was used to estimate diet diversity in insectivorous bats (Zeale *et al.* 2011). In brief, we targeted the variable region of the COX1 gene – found in all insects – to estimate insect prey diversity from gut contents of captured dragonflies. A recent study by Kamenova and colleagues (2017) showed that prey DNA remains relatively intact inside the gut of a predatory carabid beetle (*Pterostichus melanarius*) for at least 3-5 days. Assuming a similar prey retention time in dragonfly guts, we thus expected that our analysis would reflect a 3-5 day snapshot of dietary diversity in each dragonfly. We sampled a total of 45 dragonflies representing three geographical locations, three species, and two sampling seasons (Fig S1); as well as a phytophagous butterfly larva (*Hasora* sp.) as a control. We extracted dragonfly gut contents, removed the host tissue and then extracted DNA from gut contents using the Wizard ^®^ Genomic DNA Purification Kit (Promega Corporations, Wisconsin, Madison, USA) with the following modifications. We lysed cells at 65°C in nuclei lysis solution with 10 mM EDTA, followed by an overnight proteinase K treatment. We precipitated DNA overnight at -20°C, suspended the final pellet in 20 µl nuclease-free water, and checked the concentration and integrity of the DNA. For further analysis, we chose samples showing intact bands on an agarose gel (n = 28 dragonflies, and 1 butterfly larva; Table S1). We designed custom primers – containing Illumina ITS barcodes for multiplexing – to target the COX1 variable region (using references from Zeale *et al.* 2011). The forward primer sequence was 5’TCGTCGGCAGCGTCAGATGTGTATAAGAGACAGAGATATTGGAACWTTATATTTTA TTTTTGG3’, and the reverse primer was 5’GTCTCGTGGGCTCGGAGATGTGTATAAGAGACA GWACTAATCAATTWCCAAATCCTCC3’. We used 200 ng DNA from each sample to amplify the target COX1 region with High Fidelity Phusion polymerase (Thermo scientific). We purified the samples using the Qiagen PCR purification Kit (Qiagen) and checked the product for amplicon size and concentration. We prepared sequencing libraries using the Nextera XT v2 Index Kit (Illumina, U.S.A.) and sequenced them on the MiSeq platform (250 bp paired-end). Sequencing was performed by Genotypic Technology Pvt. Ltd., Bangalore, India.

We processed amplicon sequencing data using QIIME version 1.9.1 (Caporaso *et al.* 2010). After demultiplexing and removing barcodes and primer sequences, we filtered and trimmed reads for sequence length and quality score (q>20) using default QIIME parameters. We used Fast-QC to check read quality and presence of barcodes or primers in the processed data. Finally, we paired the forward and reverse reads to generate a total of 30 million high quality paired-end reads for the 16S gene, with an average of 169,000 reads per sample (range: 8,000–900,000). We classified these reads into Operational Taxonomic Units (OTUs) at the 97% similarity level using the QIIME implementation of UCLUST using closed reference (only reference based) as well as open reference (reference based and denovo) algorithms. We used the GreenGenes 16S ribosomal gene database version gg_13_8 (DeSantis *et al.* 2006) to assign taxonomy to each representative OTU. We removed chimeric sequences using Chimeraslayer (Haas *et al.* 2011) and removed unassigned, chloroplast, and mitochondrial sequences to generate the final “.biom” files for all OTU picking methods. We normalized closed referenced OTUs by bacterial 16S copy number using the software PICRUSt (Phylogenetic Investigation of Communities by Reconstruction of Unobserved States, version 1.0.0).

For insect COX1 amplicons, we obtained a total of 2.1 million reads (average 70,000 and range 29,000–115,000 reads per sample). Using QIIME, we picked OTUs at the 89% similarity level (as described in Hebert *et al.* 2003), and used the Barcode of Life Database v4 (Ratnasingham & Hebert 2007) to assign taxonomy to each OTU. We removed chimeric sequences from OTUs. We checked the precision of our sequencing and OTU assignment by examining our control sample (butterfly larva), where 97.4% of the reads were correctly classified to a single OTU assigned to *Hasora* sp without correcting for spurious OTUs. To remove host OTUs from each sample, we picked the Odonate OTU with the highest number of assigned reads and deleted it from the final table. We note that this elimination step would also remove potential cases of conspecific predation, which is known in some dragonflies (Corbet 2004).

For microbiome as well as insect diet analysis, we removed potentially erroneous OTUs (as described in Huse *et al.* 2010) from our dataset by implementing three OTU filters, generating a smaller community in each case. 1) Pruned community: retaining all OTUs with at least 0.005% relative abundance across the entire dataset, to minimize impacts of sequencing errors (Bokulich *et al.* 2013); 2) Dominant community: retaining all OTUs with at least 5% relative abundance in at least one sample; 3) Minimally pruned community: retaining all OTUs with at least 20 reads per OTU per sample, to obtain a conservative estimate of OTUs with sufficient read support. We separately applied each filter to the full dataset and then recalculated the relative abundance of OTUs for subsequent analysis.

### Statistical analysis

We performed all statistical analysis in the R statistical software version 3.3.4 (R Core Team 2013) using relevant packages as required. We considered each distinct OTU (gut bacteria: 97% sequence similarity, eukaryotic prey: 89% sequence similarity) as the basic unit of comparison, regardless of taxonomic placement. To estimate the sampling depth at which community richness saturated, we performed rarefaction analysis with the Pruned Community. We assumed that this sampling depth would be sufficient to saturate the two other pruned communities, since a) the minimally pruned community had higher reads/sample, and b) the dominant community was a subset of the pruned community. We subsampled reads to simulate varying sampling depth (100-2500 reads per sample) and calculated Faith’s phylogenetic diversity (Faith’s PD) at each depth. We plotted PD against the number of reads per sample to estimate the sampling depth at which PD saturated, as an indicator of sufficient sampling.

We analyzed community structure (relative abundance of OTUs) across samples using Ward’s hierarchical agglomerative clustering (Murtagh & Legendre 2014). We tested the impact of host species, location and season using permutational ANOVA (PERMANOVA, in the R package “Adonis” (Oksanen 2015)) with 10,000 permutations. We used the R package “Caret” (Kuhn 2008) to remove near-zero variance in the data. To visualize clustering of samples based on their bacterial composition across treatments, we calculated Bray-Curtis distances between samples and performed Canonical Analysis of Principal Coordinates based on Discriminant Analysis (CAPdiscrim) using the R package “Biodiversity R” (Kindt & Kindt 2017). We tested the significance of clustering and estimated classification success by permuting the distance matrix 1000 times. We plotted the two dominant linear discriminants (LD) to visualize data classification. For each cluster, we drew ellipses reflecting 95% confidence intervals using the function “Ordiellipse” in the R package “Vegan” (Dixon 2003; Oksanen *et al.* 2017).

To estimate bacterial or prey OTU richness for each dragonfly sample, we converted the table with the relative abundance of each OTU to a presence-absence table. We also used the final “.biom” table to identify shared OTUs across samples, and to calculate OTU richness per sample, α diversity (Shannon’s diversity index, a measure of OTU richness and evenness per sample) and β_w_ diversity (a comprehensive measure of the number of unique OTUs per sample (Koleff *et al.* 2003)). We tested the effect of host species identity, sampling location and sampling month on OTU richness using a generalized linear model (GLM) with Poisson errors.

### Quantitative PCR to validate abundance of specific gut bacteria

To estimate the abundance of eubacteria and *Wolbachia* (a prevalent insect-associated bacterial genus), we performed quantitative PCR (qPCR) on bacterial DNA extracted from 9 dragonfly guts (three individuals each of *O. sabina, O. pruinosum* and *P. flavescens*). We used previously reported primers (Heddi *et al.* 1999): universal Eubacterial primers, forward 5′-AGAGTTTGATCATGGCTCAG-3′ and reverse 5′-TACCTTGTTACGACTTCACC-3′; and *Wolbachia* specific primers, forward 5′-CGGGGGAAAAATTTATTGCT-3′, reverse 5′-AGCTGTAATACAGAAAGTAAA-3′. To normalize bacterial abundance to host tissue, we used previously described Odonate-specific primers for the 28S gene (forward: 5′-ACCATGAAAGGTGTTGGTTG-3′ and reverse: 5′-ATCTCCCTGCGAGAGGATTC-3′) (Dijkstra *et al.* 2014). All primer pairs had amplification efficiencies greater than 90%. We ran three sets of PCR for each sample (total 10 µL reaction volume), using 10 ng of host gut DNA, 8 µL SYBR green PCR master mix (Thermo Fisher Scientific, Wilmington USA), and the appropriate primers (200 nM each). We added reaction mixes in a 384 well microplate (Corning, New York, USA) and monitored amplification in a ViiA™ 7 Real-Time PCR System (Thermo Fisher Scientific, Wilmington USA) with the following cycle conditions: 95°C for 30 s, 40 cycles of 95°C for 60 s, 56°C for 60 s, 72°C for 60 s, and extension at 72°C for 5 min. We calculated threshold cycle values (C_T_) for each sample. We used the CT value of each host specific gene to estimate the ΔC_T_ values. Finally, we plotted these values for all the three host dragonflies for comparison.

### Testing models of bacterial community assembly in dragonfly guts

If the gut community of a host is under weak selection, it is expected that it will be predominantly neutrally assembled. To test whether a neutral model of community assembly could explain the observed distribution of bacterial communities across hosts, we fitted a neutral distribution model (Sloan *et al.* 2006; Woodcock *et al.* 2007) to the bacterial communities observed in hosts from a specific location and season. The model is based on Hubbell’s model of the neutral theory of biodiversity (Hubbell 2001) but is applicable to large communities, such as a complex microbial community. For model fitting, we followed the approach used by Burn et al. (2016). We considered that each individual dragonfly gut houses a local community with numerous bacterial species (OTUs) whose members are drawn from a larger metacommunity, comprised of bacteria present across all dragonfly individuals collected from a specific geographic location and season. The model uses the following parameters: (a) population size of each OTU in the local and metacommunity (estimated using the number of reads) (b) the relative abundance of each OTU. Using these, the model estimates the migration rate or dispersion probability (m) for each OTU. In the event of an individual bacterium’s death, m is the probability that it will be replaced via dispersal from the metacommunity, rather than reproduction within the local community. The relationship between the abundance of each OTU in the metacommunity and its occurrence across local communities is informative for understanding the processes driving community assembly (Sloan *et al.* 2006). Under neutral community assembly, a highly abundant OTU should occur in many hosts and fall within the 99% CI of the fitted line. If an OTU occurs at a higher frequency in a host than expected from its abundance in the metacommunity (comprised of OTUs from all hosts), this indicates positive selection for those bacteria (presumably by the host). Similarly, if an OTU is very abundant in the metacommunity but occurs in only a few host individuals, this indicates negative selection against the OTU.

For each metacommunity derived from hosts sampled from a given location in a specific season, we fitted a β-distribution to the relationship between OTU occurrence and abundance (using the script published by Burns et. al. 2016). We checked the model fit using non-linear least squares in R, and estimated 99% confidence intervals (CI) around the fit using binomial proportions. We then compared the proportion of OTUs that were neutrally distributed across sites, seasons and hosts. We generated two sets of models: (1) for each dragonfly species sampled at a specific location and season (2) pooling all dragonfly species sampled in each location and season. The first set allowed us to infer patterns of gut bacterial community assembly for each dragonfly species; but with low sample sizes (Table S1). The second set allowed us to infer general patterns of gut bacterial assembly across dragonflies, with a larger sample size. Finally, we compared taxonomic diversity (Clarke & Warwick 1998; Fierer *et al.* 2007; Morrow *et al.* 2015) between the groups of bacteria that were inferred to be neutrally distributed, or positively or negatively selected by hosts. If the hosts were selecting for a specific functional association (and this functionality is phylogenetically conserved in bacteria), we expected that bacterial OTUs experiencing positive host selection should have lower taxonomic diversity compared to neutrally assembled bacteria.

### Localizing bacteria in dragonfly guts

We used fluorescent in-situ hybridization (FISH) to determine the location of bacteria inside dragonfly guts. We hypothesized that if there is a functional association between host and bacteria, bacterial cells should be housed in specific crypts or inside columnar cellular folds in the host gut (as reported in previous studies by Barrow *et al.* 1980; Fuller & Turvey 1971). However, if bacteria are transient or only associated with food particles, they should be easily washed off or would be found primarily in the inner lumen of the host gut. We performed FISH with three species of dragonflies (*O. sabina, O. pruinosum,* and *P. flavescens*; n = 5 per species), using bacteria-specific probes (universal eubacterial probe, [Alexa-488] 5’-GCTGCCTCCCGTAGGAGT-3’ (Da Silva *et al.* 2015) and a *Wolbachia* specific probe, [ALEXA-647] 5’-CTTCTGTGAGTACCGTCATTATC-3’ (Le Clec’h *et al.* 2013), obtained from Sigma-Aldrich-Merck, Missouri, USA) and DAPI (4’, 6-diamidino-2-phenylindole) staining (to visualize host cell nuclei). We followed the COLOSS protocol (http://www.coloss.org/2018/01/17/standard-methods-for-molecular-research-in-apis-mellifera/) with a few modifications. Before the assay, we rehydrated guts and fixed them in Carnoy’s fixative for 96 hours. To reduce autofluorescence, we used peroxide treatment for 72 hours and replaced the water inside host tissues using absolute alcohol and xylene washes as per the COLOSS protocol. Finally, we embedded samples in liquid paraffin using plastic molds to create paraffin blocks. We sliced the blocks into 10 μm transverse sections using a Leica manual microtome (Leica Microtome 2125 RTS, Wetzlar, Germany) using disposable blades (Low profile blade 819, Leica). For each species and each part of the gut (foregut, midgut and hindgut), we obtained 5 sections per probe for each of 5 individuals. We mounted sections on Fisherbrand Superfrost Plus microscope slides (Thermo Fisher Scientific, Wilmington, USA), and heated off the paraffin in an oven at 65°C. We washed with Xylene (three minute wash, thrice), absolute alcohol (three minute wash, thrice), and double-distilled water (once) before hybridization. We dissolved 0.5µL of fluorescent probes in 500µL hybridization buffer and stained gut tissue sections in a dark chamber for 8-10 hours at room temperature with the respective bacteria-specific probe. We then stained sections with DAPI for 20 min to visualize host cell nuclei. We applied DABCO-glycerol (antifade agent), sealed the sections with coverslips, and stored them at 4ºC in the dark. We note that we lost many sections during the multiple washes, but we retained at least 2 sections/dragonfly/species/probe for final analysis. We imaged sections using a Zeiss 510 Meta confocal microscope (Carl Zeiss, Oberkochen, Germany). We analyzed images using Image-J software (Version 1.6.0-24, 64-bit version).

## 3. RESULTS

Initial rarefaction analysis revealed that our sampling depth was sufficient to determine the bacterial community composition in all but one sample, which we excluded from further analysis (Fig S1). We separately analyzed a total of six sets of bacterial communities, generated using either closed or open-reference OTU picking and implementing three OTU filtering thresholds: 1) pruned community (576 OTUs open reference), 2) dominant community (59 OTUs open reference), and 3) minimally pruned community. All sets showed comparable results, but here we focus on the pruned and dominant open referenced sets unless mentioned otherwise. Corresponding results for other sets are given in the supplementary material.

We found an average of 188 OTUs per sample, which is over 2-fold higher than previous observations for other carnivorous insects including the order Odonata (Ley *et al.* 2008; Sullam *et al.* 2012; Jones *et al.* 2013; Yun *et al.* 2014). The dragonfly gut community was dominated by Proteobacteria (88%), Firmicutes (9.8%), Actinobacteria (1.8%) and Bacteroidetes (0.4%) (Figs 1B & 1C). At the family level, Rickettsiaceae – comprising of three *Wolbachia* OTUs – were most abundant, although this high abundance was limited to dragonflies from the genus *Orthetrum* (Fig 1D). In other host genera, especially *P. flavescens*, OTUs from the family Enterobacteriaceae were more abundant. Overall, we observed substantial variation in relative abundance of OTUs across individual hosts (Fig 1D).

### Host species, sampling season and location shape gut bacterial community composition

A full analysis of the impact of host species, season and location showed that each of these factors had significant impacts on the dominant gut bacterial community (Table 1; see Table S2 for other community sets). Linear discriminant analysis to visualize clustering supported the PERMANOVA analysis, showing strong separation in gut bacterial communities across host species, location, and season (Fig 2, and S2). An unconstrained PCoA analysis (Figs S3) also showed similar patterns, although as expected, classification ability was poorer than observed for constrained (CAPdiscrim) analysis (Fig 2). Interestingly, location and season together explained a larger proportion of variation in gut bacterial communities (24%; Table 1A) compared to host species alone (16%), suggesting that environmental factors might have a stronger impact on community composition. In fact, when we restricted our analysis to the three best-sampled host species, *O. sabina, O. pruinosum,* and *P. flavescens*, we found a relatively weak impact of host species (7% variation explained), relative to the combined effects of location and season (total 31% variation explained; 22% by location alone) (Table 1B, and S3). These patterns are also mirrored in the number of shared vs. unique bacterial taxa across various groups of dragonflies. Out of the 576 OTUs detected in total, all host species shared 206 OTUs (~36%; Fig S4G). Interestingly, the congeneric dragonflies *O. pruinosum* and *O. sabina,* which harbored similar bacterial communities (Fig 2A), also shared the maximum number of bacterial OTUs (407 shared OTUs, ~71%, out of which 34 OTUs were unique to the genus *Orthetrum*; Fig S4G). Finally, 42% of the OTUs (241 out of 576) were shared across seasons (Fig S4H), and 25% (145 out of 576 OTUs) were shared across locations (Fig S4I).

**Figure 2.**
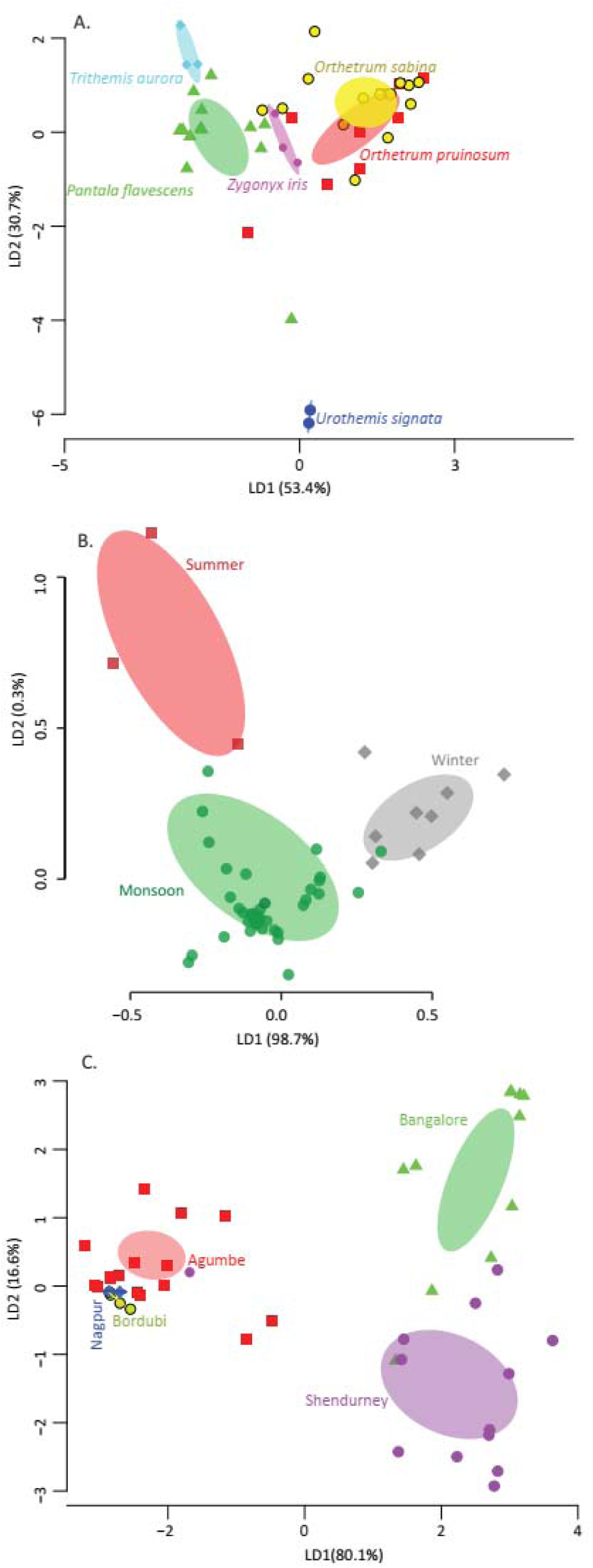
Linear discriminant (LD) plots showing two dominant linear discriminants (LD) that group dragonfly samples based on their gut bacterial community composition (based on Bray-Curtis distance and open reference OTU picking). Percentage of variance explained by each LD is indicated in parentheses. Each point represents a host individual. Ellipsoids represent 95% confidence intervals around each group mean, calculated from LD values. Clustering of dragonfly samples based on (A) host species identity (B) sampling season (C) sampling location.

**Table 1.** Results of a permutational analysis of variation (PERMANOVA) showing the effect of host species, location, and sampling season on gut bacterial composition (open referenced dominant bacterial community) (A) across all six dragonfly hosts, and (B) across the three best-sampled dragonfly hosts (*Orthetrum pruinosum*, *Orthetrum sabina*, and *Pantala flavescens*).

| (A) | Df | SSq. | Mean SSq. | F stat | R<sup>2</sup> | P |
| --- | --- | --- | --- | --- | --- | --- |
| Species | 5 | 3.01 | 0.60 | 1.92 | 0.16 | 0.0004 |
| Location | 4 | 2.81 | 0.70 | 2.24 | 0.15 | 0.0003 |
| Season | 2 | 1.83 | 0.91 | 2.91 | 0.09 | 9.999e-05 |
| Interaction (Species, Location) | 3 | 1.64 | 0.55 | 1.74 | 0.08 | 0.006 |
| Residuals | 32 | 10.05 | 0.31 |  | 0.52 |  |
| Total | 46 | 19.34 |  |  | 1 |  |
| (B) | Df | SSq. | Mean SSq. | F stat | R<sup>2</sup> | P |
| Species | 2 | 0.98 | 0.49 | 1.76 | 0.07 | 0.042 |
| Location | 4 | 3.20 | 0.80 | 2.88 | 0.22 | 4e-05 |
| Season | 2 | 1.41 | 0.70 | 2.54 | 0.09 | 0.003 |
| Interaction (Species, Location) | 3 | 1.42 | 0.47 | 1.70 | 0.10 | 0.03 |
| Residuals | 27 | 7.51 | 0.28 |  | 0.52 |  |
| Total | 38 | 14.5 |  |  | 1 |  |

Overall, the impacts of host species, location and season were not sensitive to omission of the two most abundant bacterial families Rickettsiaceae and Enterobacteriaceae (Figs S5A-B), indicating a robust bacterial community structure. Conversely, focusing only on these two bacterial families, we found slightly different results. The abundance of OTUs from the family Rickettsiaceae was influenced by both host species and sampling location (Table S4A), with high abundance in the genus *Orthetrum* (average 82%, Figs 1D and S6A; confirmed using qPCR, Fig S7) except for Bangalore samples (<1%) (Fig S6B). The abundance of Enterobacteriaceae OTUs was influenced by host species (predominant in *P. flavescens*, average 70%) and sampling season (predominant during monsoon, average 42%) (Figs 1D and S6C-D; Table S4B). Finally, for each factor, classification analysis based on gut bacterial composition also categorized significant proportions of samples correctly into the respective groups (Tables S5A-C and S6A-C). These results show that host-specific and environmental factors together govern bacterial community structure, with the latter having larger impacts.

Despite the significant effects of host species, location and season on overall community composition, these factors had relatively weak and variable impacts on the richness and diversity of bacterial communities. Across host species, location and season, bacterial communities had similar number of OTUs (Fig S8 A-C; Table S7A). Similarly, the α diversity of communities (considering both OTU richness and evenness) varied only across host species (Fig S4A; Table S7B, and S8), but was invariable across sampling season and sites (Figs S4B-C; Table S7C). In contrast, all three factors (as well as an interaction between location and host species) significantly affected the β diversity of communities (Figs S4D-F; Table S7C), indicating significant community turnover across species, season and site. However, the impact of host species on β diversity was largely driven by the two *Orthetrum* species and *P. flavescens*, all of which had higher β diversity than the other three hosts (Fig S4D). Interestingly, dragonflies collected during the monsoons also showed greater β diversity (Fig S4E), suggesting an impact of rainfall on gut bacterial diversity. β diversity was also higher in sites from Southern India (Agumbe, Bangalore and Shendurney) compared to North Indian locations (Bordubi and Nagpur) (Figs S4F). However, the reasons for these differences in β diversity are not obvious: Agumbe, Shendurney and Bordubi are close to rainforests with relatively high biodiversity, whereas Nagpur and Bangalore are dry areas with relatively low biodiversity.

### Dragonflies show host-specific and seasonal dietary specialization

To test whether host-specific bacterial communities reflect host-specific diets, we next tested for dietary specialization across the three best-sampled dragonfly species, *O. pruinosum, O. sabina* and *P. flavescens.* After excluding putative host OTUs, the pruned prey community of both *Orthetrum* species had significantly higher richness compared to *P. flavescens* (P<0.01, Chi-square=16.91, df=2, post-hoc Dunn test: OP vs. PF: P<0.01, OS vs. PF: P<0.01, OP vs OS: P=0.04) as well as greater diversity (P=0.01 Kruskal Wallis’ Chi-squared: 8.37, df=2, post-hoc Dunn test: OP vs. PF: P=0.03, OS vs. PF: P<0.01, OP vs OS: P=0.26) (Fig 3A). These patterns mirror the bacterial communities associated with these hosts (Fig 3A):*Orthetrum* had higher bacterial diversity (Kruskal Wallis’ Chi-squared: 7.39, df=2, P=0.02, post-hoc Dunn test: OP vs. PF: P=0.03, OS vs. PF: P=0.04, OP vs OS: P=0.69), though not significantly higher richness (Kruskal Wallis’ Chi-squared: 1.99, df=2, P=0.3). These correlated differences in prey and gut bacterial communities indicate a potential link between the two.

**Figure 3.**
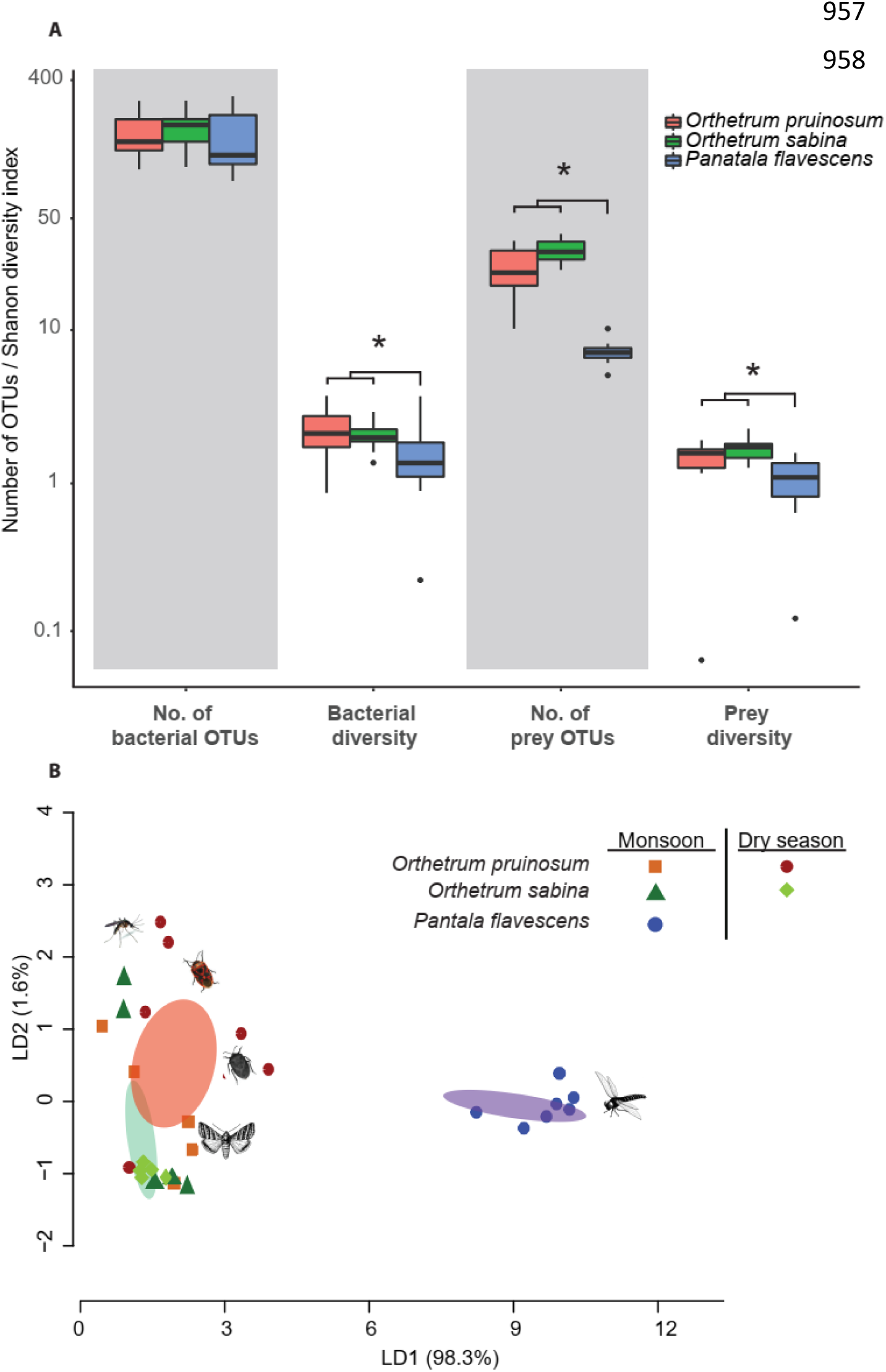
(A) Boxplots show OTU richness and diversity of bacterial and prey communities of three dragonfly species (*Orthetrum pruinosum, Orthetrum sabina* and *Pantala flavescens*). Asterisks indicate significant differences in OTU diversity or richness (Kruskal Wallis test). (B) Clustering of dragonfly samples based on dietary composition using LD analysis, as described in Fig 2. Representative images of dominant prey taxa are shown for each dragonfly species.

We also observed striking differences between the diets of the three dragonfly species, with the two congeneric *Orthetrum* species sharing more similar diets (Fig 3B, and S9, Table 2, Table S10). For *O. pruinosum* and *O. sabina,* the prey community was predominantly composed of Dipterans (83% and 68% respectively) (Fig S9), whereas *P. flavescens* consumed more Odonates (88% of prey OTUs) (Fig S9). We also observed that the diets of the two *Orthetrum* species changed across seasons (Table 2; Figs 3B and S10, Table S9). During the monsoon, individuals of both species had similar diets, but during the dry season, they had more dissimilar diets (Fig 3B and S10, Table S9). For *O. pruinosum,* diet α-diversity and richness tended to increase during the monsoon (diversity: Welch t-test: t=-1.71, df=7.31, P=0.06; richness: Welch t-test: t=-2.4, df=7.66, P=0.01) (Fig S10), potentially due to increased availability of diverse insect prey after the rains. However, the diet diversity of *O. sabina* reduced during the monsoon (Mann-Whitney U test: W=21, P=0.04, Fig S10), without any impact on dietary richness (Welch t-test: t=-0.05, df=7.9, P=0.5). We found that during monsoon, diet evenness also decreased in *O. sabina* (Welch t-test: t=-1.71, df=5.48, P=0.07, marginally non-significant), whereas it remained unaltered in *O. pruinosum* (Welch t-test: t=-0.8, df=6.01, P=0.22) (Fig S10). Note that we did not observe strong impacts of location on dragonfly diet (Table 2; no effect of location alone, and a marginally significant effect of interaction with host species). This potentially reflects our limited sampling: we could only sample from two very closely located sites (Agumbe and Kadhari), which have similar habitats and probably similar insect prey communities. Overall, these results suggest that (a) *O. sabina* is likely a specialized forager whose preferred prey are more abundant during monsoon (thus decreasing evenness and diversity without affecting richness) (b) *O. pruinosum* is a generalist predator whose prey base diversifies depending on prey availability and (c) *P. flavescens* is a specialized predator that predominantly targets other Odonates (Figs 3B, S9, and S10). These patterns are also consistent with the hypothesis that host- and season-specific gut bacterial communities of dragonflies may directly reflect the influence of the introduction of diet-specific bacteria into the insects’ guts.

**Table 2.** Results of a permutational analysis of variation (PERMANOVA) showing the effect of host species, location, and season on the diet of *Orthetrum pruinosum*, *Orthetrum sabina*, and *Pantala flavescens.*

|  | Df | SSq. | Mean SSq. | F stat | R<sup>2</sup> | P |
| --- | --- | --- | --- | --- | --- | --- |
| Species | 2 | 2.97 | 1.48 | 9.53 | 0.36 | 9.00E-05 |
| Location | 1 | 0.21 | 0.21 | 1.38 | 0.027 | 0.2 |
| Season | 2 | 1.15 | 0.57 | 3.69 | 0.14 | 4.00E-04 |
| Interaction (Species, season) | 1 | 0.44 | 0.44 | 2.85 | 0.05 | 0.021 |
| Interaction (Species, location) | 1 | 0.31 | 0.31 | 2.03 | 0.04 | 0.056 |
| Residuals | 20 | 3.11 | 0.155 |  | 0.37 |  |
| Total | 27 | 8.21 |  |  | 1 |  |

### Dragonfly gut bacterial communities are predominantly neutrally assembled

To specifically test the hypothesis that dragonfly gut bacterial communities are acquired passively through the diet – with relatively weak host imposed filters – we estimated the fraction of the bacterial community whose occurrence and abundance across hosts was consistent with neutral vs. non-neutral assembly. Analyzing communities from all samples collected from a given location and season (regardless of host species), we found that a large fraction of bacterial OTUs are predicted to be neutrally assembled (mean 76 ± 0.09 %; range; Figs 4A-B, and S11A-C); i.e. whose distribution across hosts matched expectations from a model simulating assembly via random OTU dispersal. The proportion of neutrally distributed gut bacteria was influenced by both location and sampling season, with no interaction between these two factors (Table 3A). Dragonflies from Bordubi had the highest proportion of neutrally assembled gut bacteria (83%; Figs 4A and S12C), whereas dragonflies from Nagpur had the lowest proportion (62%; Figs 4A and S12E). Hosts collected in the dry season had a higher proportion of neutrally assembled gut bacteria (84%) compared to the monsoon (71%) (Fig 4B), potentially reflecting fewer dietary options available during the dry season. This pattern was also reflected in Agumbe which was sampled comprehensively across both seasons (proportion of neutrally assembled community: monsoon: 67%, dry season: 85%). As expected, we observed reverse patterns for the relative fraction of OTUs whose distribution is consistent with positive selection (Table 3B, Figs 4C-D) or negative selection (Table 3C, Figs 4E-F). Interestingly, monsoon samples had higher proportions of bacteria under negative selection (Fig 4F), suggesting that despite a high influx of new bacteria during monsoon, many of these could not establish in the hosts, and had low abundance. Finally, pooling all OTUs predicted to be under positive selection (across locations and seasons), we found that the taxonomic diversity of these was either higher than or comparable to OTUs that were neutrally distributed or under negative selection (Fig S13). This result is inconsistent with the hypothesis that dragonflies impose strong positive selection favoring a specific, shared set of functionally important bacteria. Instead, the pattern of higher taxonomic diversity in positively selected bacteria is consistent with host-specific, seasonal or spatial variation in putatively beneficial bacteria.

**Figure 4.**
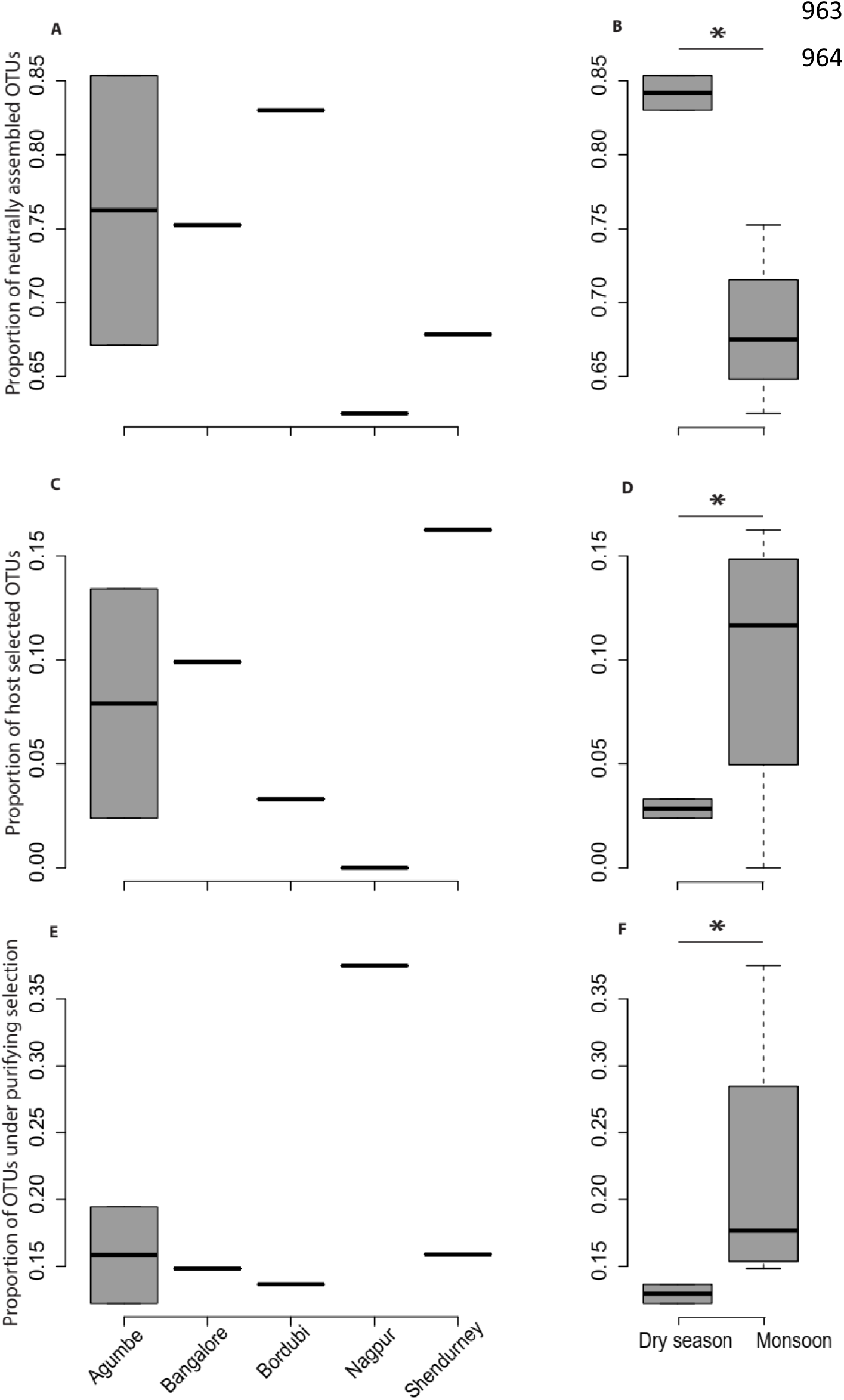
Boxplots show the proportion of bacteria whose distribution is consistent with (A, B) neutral assembly (C, D) positive selection and (E, F) negative selection, for dragonflies sampled from a given location or in a specific season. In panels B, D and F, asterisks indicate a significant difference across seasons.

**Table 3.** Results of GLMs (Generalized linear models) testing the effect of location and season on the proportion of dragonfly gut bacterial community that is (A) neutrally assembled (B) under positive selection and (C) under purifying selection.

| (A) | Df | Deviance | Residual deviance | Deviance | P |
| --- | --- | --- | --- | --- | --- |
| Null |  |  | 5 | 52.5 |  |
| Location | 4 | 27.07 | 1 | 25.5 | 1.919e-05 |
| Season | 1 | 25.5 | 0 | 0 | 4.370e-07 |
| (B) | Df | Deviance | Residual deviance | Deviance | P |
| Null |  |  | 5 | 49.68 |  |
| Location | 4 | 44.33 | 1 | 5.34 | 5.476e-09 |
| Season | 1 | 5.34 | 0 | 0 | 0.03 |
| (C) | Df | Deviance | Residual deviance | Deviance | P |
| Null |  |  | 5 | 83.8 |  |
| Location | 4 | 59.1 | 1 | 24.7 | 4.49e-12 |
| Season | 1 | 24.7 | 0 | 0 | 6.67e-07 |

Since we had relatively low sample sizes for each dragonfly species in a given location and season (n=3), we restricted our analysis (above) mainly to pooled results across all samples. However, we also attempted to investigate host species-specific patterns of gut bacterial community assembly (Fig S11). We found that both host species and location had a significant impact on the proportion of bacteria that are neutrally assembled (P<0.01 in each case; Table S11A; Fig S11), but that season had no effect (P=0.2, Table S11A) (also see Table S11B for bacteria under positive selection). Interestingly, for the two *Orthetrum* species, the pattern of gut bacterial assembly across sampling seasons was concordant with the change in their diets. We found that the proportion of neutrally assembled gut bacteria in *O. pruinosum* – a generalist predator – was higher during the monsoon (mean: 79%, median: 84%) than in dry season (mean & median: 70%) (Fig S14A). On the contrary, *O. sabina* – a relative specialist during the monsoon – showed an inverse pattern (monsoon: mean 70%, median 62%, dry: mean & median 77%; Figs S14A and B). However, it is important to be noted that we could not perform relevant statistical tests due to insufficient sample size. These patterns support the hypothesis that diet-driven neutral community assembly likely explains much of the variation that we observe in gut bacterial community structure in dragonflies.

### Bacterial cells rarely adhere inside dragonfly guts

To test whether bacterial cells adhere to dragonfly guts or are housed in specialized structures, we dissected the guts of three species (*O. sabina, O. pruinosum,* and *P. flavescens*) and probed for bacteria using FISH (Fig 5). The gut lumen was lined with columnar folds of epithelial cells (Fig 5A); in case of a specific host-bacterial association, bacteria could adhere or be housed here. However, we did not find any eubacterial signal in the foregut (Fig 5B-D), indicating that bacteria were either absent or rare in this part of the gut. Since we did not find a signal with the general eubacterial probe, we did not test foregut sections with the *Wolbachia*-specific probe. In *P. flavescens*, only the eubacterial probe showed a positive signal inside columnar folds (5 of 5 tested individuals; 3 with very small patches of bacteria) (Fig 5E and H), whereas *Wolbachia* was absent (Fig 5K and N), corroborating our amplicon sequencing results. The midgut and hindgut of both *Orthetrum* species were positive for eubacterial and *Wolbachia*-specific probes (Fig 5F, G, I, J, L, M, O, and P; all 5 tested individuals of each species), although the signal was generally weak and localized to a small cluster of bacteria found in the gaps between columnar cellular folds. Interesting exceptions were observed in two *O. sabina* individuals where *Wolbachia* appeared to be sequestered within a specific tissue structure (Fig 5L); the functional significance of this pattern requires further work. Overall, the lack of a predominant signal of gut colonization suggests at best a weak relationship with the host.

**Figure 5.**
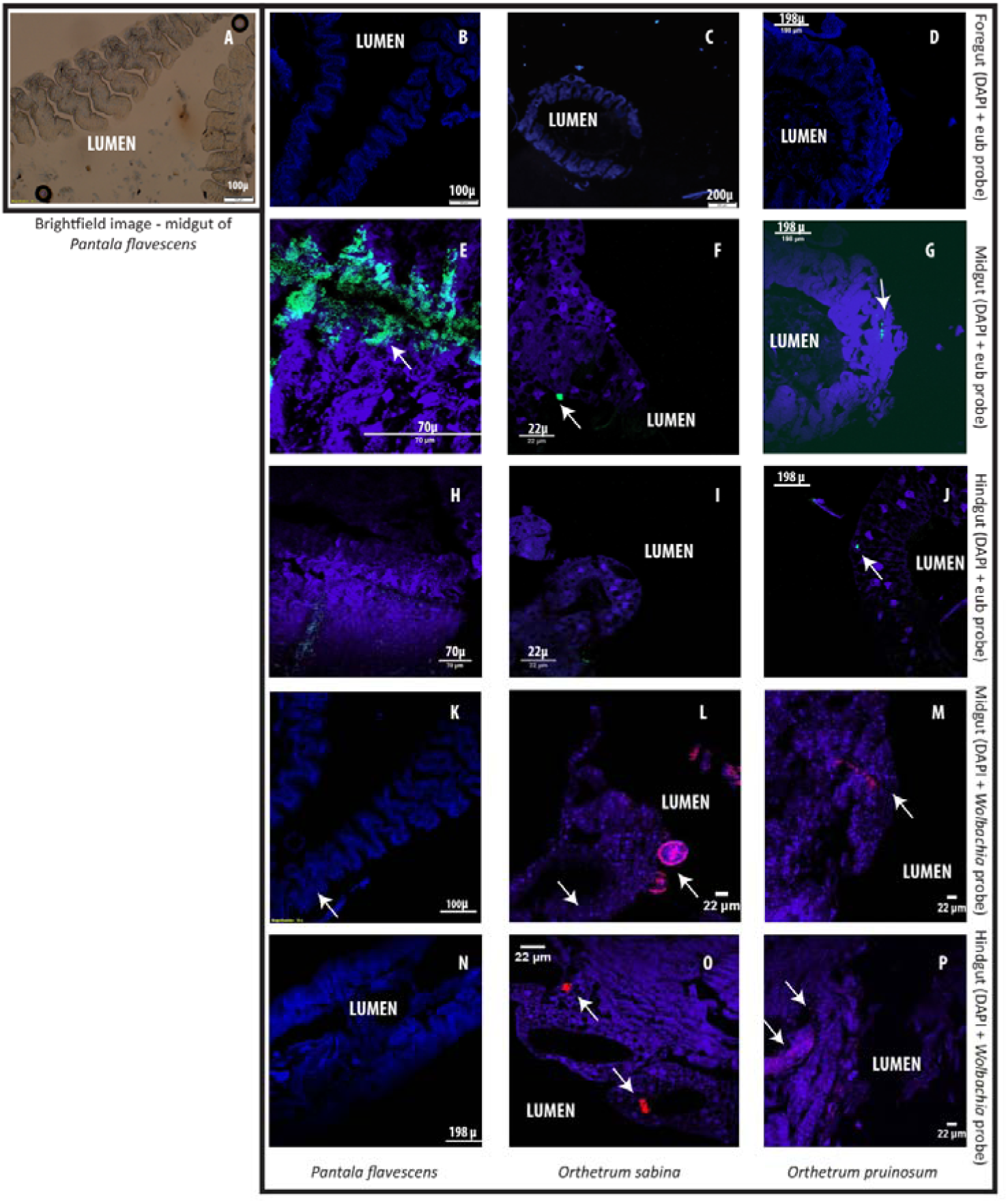
Examples of Fluorescent in situ hybridization (FISH) images of dragonfly gut sections using bacteria-specific probes. Host cell nuclei are stained purple with DAPI; eubacteria are green; and *Wolbachia* are pink. Arrows highlight bacteria in each section. (A) Representative brightfield image of *P. flavescens* midgut section showing columnar cellular folds covering the gut lumen, and food particles in the lumen. (B-D) Foregut sections of *P. flavescens*, *O. sabina* and *O. pruinosum.* Note the lack of eubacterial or *Wolbachia* signal. (E-G) Midgut and (H-J) hindgut sections of each species, stained with a eubacterial probe. Note the strong eubacterial signal near the columnar folds of *P. flavescens*. (K-M) Midgut and (N-P) hindgut sections of each species, stained with a *Wolbachia-* specific probe. Note the lack of signal in *P. flavescens,* a weak signal in *O. pruinosum*, and a large globular structure with *Wolbachia* in *O. sabina.*

## 4. DISCUSSION

Host selection is generally considered to be a strong selective force shaping gut bacterial communities of animals (Colman *et al.* 2012; Engel & Moran 2013; Yun *et al.* 2014), and is expected to stabilize communities in the face of spatial and temporal variation. Here, we tested this prediction by analyzing host associated gut bacteria across spatially and temporally separated populations of six dragonfly species. Our key results contrast multiple findings from prior work: (a) dragonfly bacterial communities are twice as rich and diverse as other carnivorous insects including Odonates (Ley *et al.* 2008; Sullam *et al.* 2012; Jones *et al.* 2013; Yun *et al.* 2014); (b) location and season together explain more variation in bacterial community composition than host species identity; (c) dragonflies have somewhat specialized diets that reflect patterns of variation in gut bacterial communities, and (d) the gut community is predominantly neutrally assembled, showing little signatures of the strong host selection reported for many other insects (Engel & Moran 2013; Yun *et al.* 2014). Thus, our work highlights the importance of analyzing gut microbial communities of natural host populations in the context of naturally observed variation in geography, season, and host taxonomy.

In conjunction with host-imposed selection, ecological variables such as geographic and seasonal variation are hypothesized to contribute to the assembly of unique gut bacterial communities across insect hosts (Hufeldt *et al.* 2010; Osei-Poku *et al.* 2012; Engel & Moran 2013). Geographical variation is thought to be important because local bacterial diversity, environmental conditions, and host diet may all vary in space, and can potentially influence bacterial community composition (Dillon & Dillon 2004; Osei-Poku *et al.* 2012; Engel & Moran 2013). For instance, a specific subset of environmental microbiota associated with diet can enter host guts (dispersal), and can change either passively (through drift-like processes) or through active selection (imposed by host immunity, within host environment, or other microbial taxa in the gut). In the absence of strong host selection for specific microbes, neutral processes may dominate and lead to distinct community structure across geographically isolated host populations. For instance, previous studies found significant geographical structure in well-studied species such as humans, flies, and bees (Corby-Harris *et al.* 2007; Turnbaugh *et al.* 2009; Costello *et al.* 2009). Concordantly, our work reveals that the dragonfly gut bacterial community varies significantly across locations. Importantly, we find that community structure is not obviously affected by the geographical distance between sites. For instance, dragonflies of the same species collected from relatively close sites (Fig 1A) – Bangalore, Agumbe, and Shendurney – had distinct gut bacterial community composition, suggesting that a combination of multiple locally acting factors may drive the composition of site-specific gut bacterial communities. These factors may include specific environmental conditions (e.g. temperature, precipitation, and soil pH) that drive variation in environmental microbes; variation in insect prey communities driving differential dispersal into host guts; local host diet specialization; or site-specific variation in host imposed selection acting on similar environmental microbes.

Apart from geographical variation, seasonal variation can also influence gut microbial community by altering the environmental bacterial composition, host physiology, and host diet (Dillon & Dillon 2004; Engel & Moran 2013). We found that dragonflies housed unique gut bacterial communities across seasons, showing higher β diversity during monsoon. This impact of rainfall may occur because rain can alter the presence or abundance of environmental bacteria and/or prey species. Although the impact of climatic shifts on microbes is debated (Fierer *et al.* 2007; Jones *et al.* 2013; Wei *et al.* 2014), it is likely that increased humidity in the monsoon is more conducive for microbial growth. Prior work shows that insect diversity (major prey of dragonflies) also responds rapidly to change in rainfall (Tauber *et al.* 1986; Pinheiro *et al.* 2002; Akorli *et al.* 2016), especially in the tropics. Since insects house host-specific microbiota (Dillon & Dillon 2004; Engel & Moran 2013; Yun *et al.* 2014), a shift in the prey base may thus directly or indirectly contribute to changes in the gut bacterial community of dragonflies. Because our sampling across seasons was limited, our results probably present a conservative estimate of seasonal variation in gut bacterial communities of dragonflies.

Finally, our study revealed that each dragonfly host genus housed a distinct gut bacterial community irrespective of sampling season and location. Interestingly, both species from the genus *Orthetrum* shared a significant proportion of their gut community, which may suggest a role for phylogenetically conserved host level processes shaping the gut community. Host taxonomy is an important factor that structures gut microbiota through active or passive filters imposed by host morphology, physiology, development, immune function, social interactions or diet (Dillon & Dillon 2004; Sullam *et al.* 2012; Colman *et al.* 2012; Jones *et al.* 2013; Engel & Moran 2013; Aksoy *et al.* 2014; Yun *et al.* 2014). However, in our analysis, host taxonomy explained less variation in bacterial communities, compared to the combined impacts of location and season. For instance, despite significant host specificity, all dragonflies shared a substantial proportion (36%) of bacterial OTUs. Hence it was important to ask: what processes explain these patterns of partially shared components of gut microbiomes?

Broadly speaking, host-specific gut microbiota may reflect host specific diets and/or host specific selective filters (Colman *et al.* 2012; Engel & Moran 2013). Unfortunately, information on dragonfly diet is scarce, because their rapid and unpredictable movement patterns make observations very difficult (Corbet 2004). Limited behavioral observations in natural and cultured populations suggest that dragonflies are generalists (Fraser 1933; Corbet 2004; Stoks & Córdoba-Aguilar 2012). We used a molecular approach to identify recent insect prey in dragonfly guts, thus presenting the first precise understanding of dragonfly diet in natural conditions. Our results revealed that three common, sympatric dragonflies (*O. sabina*, *O. pruinosum*, and *P. flavescens*) consume distinct insect communities. Importantly, this dietary specialization was also reflected in their gut bacterial community composition; although we caution that our analysis shows a correlation, but not direct causation. The two *Orthetrum* species showed a degree of dietary overlap, comparable to the overlap in their gut bacterial communities; whereas *P. flavescens* was unique both with respect to its diet and its gut bacterial community. Both *Orthetrum* species had a diverse prey base with ~40 OTUs, most belonging to the order Diptera. In contrast, *P. flavescens* consumed less than 10 OTUs that were predominantly comprised of Odonates. These findings were supported by behavioral observational data (Fraser 1933; Corbet 2004) and our own unpublished data, where *Orthetrum spp.* were observed to prey on flies and mosquitoes and *P. flavescens* was found to predate on other dragonflies. These dietary differences were also reflected in the diversity and richness of gut bacteria, strongly suggesting a direct association between dietary and gut bacterial diversity. Similar patterns have been documented in other insects where diet plays a key role in gut microbe composition (Colman *et al.* 2012; Engel & Moran 2013).

Our results for the two *Orthetrum* species suggest that dietary specialization may also explain seasonal variation in dragonfly bacterial communities. The dietary overlap between these species arose primarily from similar diets during the rains; whereas their diet differed during the dry season. We speculate that *O. pruinosum* is a more generalist species, consuming more diverse and abundant prey as they become available during the monsoon, leaving prey community evenness unperturbed. On the other hand, *O. sabina* appears to be a seasonal specialist, consuming a smaller subset of more abundant prey during monsoons, leading to a decrease in prey diversity and evenness but maintaining similar prey richness as in the dry season. Thus, each dragonfly species may have a unique dietary niche that acts as a passive filter modulating the entry of environmental microbiota into the gut. Although this hypothesis requires further validation, we suggest that such dietary specialization – rather than strong host selection – is the primary driver of variation in dragonfly gut bacterial communities. Indeed, simulations using Sloan’s neutral assembly model (2006) revealed that bacterial communities were predominantly neutrally assembled. Although such modeling has only rarely been used, its application to laboratory-reared zebrafishes (Burns *et al.* 2016) and wild *Drosophila* populations (Adair *et al.* 2018) revealed largely neutrally assembled gut microbiota. In our analysis, the gut bacterial assembly also varied across location and season, with stronger signatures of neutral assembly during the dry season (summer and winter). This is expected in relative dietary specialists such as *O. sabina,* where limited prey options during the dry season should reduce specialization, leading to stronger effects of neutral processes. Thus, in the monsoon, the probability of bacterial dispersal from the metacommunity to the local community would decrease, making the local gut community of *O. sabina* more distinct from the larger metacommunity. In contrast, for a generalist host such as *O. pruinosum*, the proportion of neutrally assembled gut bacteria increased with increasing prey diversity during monsoon.

Finally, two other lines of evidence support our conclusion that dragonfly gut bacterial communities are largely structured via passive processes. First, the high taxonomic diversity of “selected” bacterial OTUs suggest the lack of preference for a specific set of phylogenetically conserved functional traits. Second, FISH with eubacterial probes failed to show strong signatures of bacteria adhering to gut walls or enclosed in specific structures, indicating a weak association with hosts. Interestingly, we observed two cases where *Wolbachia* was found inside globular sacks or crypts in *O. sabina* individuals. Such structures – thought to provide a conducive environment for bacterial proliferation – were previously reported in insect hosts with a strong association with specific microbes (Dillon & Dillon 2004; Engel & Moran 2013). Prior studies examining the association between *Wolbachia* and *Drosophila* host found that the presence of *Wolbachia* prevented further infection in hosts (Hedges *et al.* 2008; Osborne *et al.* 2012). Thus, the association between *O. sabina* and *Wolbachia* suggested by our analysis deserves further attention as a possible special case of strong dragonfly-bacterial interactions.

## Conclusions

Our analysis of patterns of spatial, temporal and host-specific variation in the diet and gut bacterial communities of multiple wild-collected dragonflies highlights two key points. First, we suggest that environmental factors that may alter bacterial community stability should be given more importance while drawing general conclusions about host-microbe interactions. Second, while explaining variation in microbial community composition, it is important to explicitly consider neutral processes along with selection. We acknowledge that our sampling effort to understand gut bacterial diversity across seasons and locations was limited. However, despite this limitation, we found considerable variation (across season and location) in our study which is likely to increase with greater sampling effort. Hence our study provides a conservative estimate of the natural variation present across populations of predatory dragonflies. Moreover, in our subsampled data with the three well sampled dragonflies we found consistent pattern. We hope that our work encourages further analysis of variation in gut microbiomes of natural insect populations, as well as experimental tests of the role of neutral vs. selective processes in the assembly of host-associated microbial communities.

## DATA ACCESSIBILITY

All data and custom code will be made available in public repositories. Sequencing data and metadata will be available on the ENA website. OTU tables (.txt files) will be uploaded in Figshare. Custom R scripts for analysis will be available on GitHub.

## ACKNOWLEDGMENTS

We thank Agashe lab members for critically reading the manuscript. We thank Krushnamegh Kunte for contributing samples from Shendurney Wildlife Sanctuary (permit no. WL 10-3781/2012 dated 18/12/2012, and GO (RT) No. 376/2012/F and WLD dated 26/07/2012); Agumbe Rainforest Research Station, Rohini Balakrishnan, and Sarita and Ramanuj Dasgupta for logistical support; Ashish Tiple, Krushnamegh Kunte, Kruttika Phalnikar, Manjunatha Reddy, Ronita Mukherjee, Sudhakar Gowda and Saira Guha for field assistance; Shantanu Joshi for help with identifying dragonflies; and Kruttika Phalnikar for help with QIIME analysis. We acknowledge funding and support from the National Centre for Biological Sciences and the Department of Science and Technology, India (INSPIRE Faculty award IFA-13 LSBM-64 to DA).

## AUTHOR CONTRIBUTIONS

RD: analyzed Miseq data; designed work, collected and prepared samples, and analyzed data for diet, FISH and qPCR experiments; carried out community assembly analysis; prepared figures and drafted the manuscript. AN: collected and prepared samples for gut bacterial analysis. DA: conceived the study; designed experiments; collected samples; acquired funding; wrote the manuscript.

